# Molecular evolution and adaptation of SARS-CoV-2 omicron XBB sub-lineage Spike protein under African selection pressure

**DOI:** 10.1101/2023.08.16.553557

**Authors:** Milton S Kambarami, Ngorima Godwins, Praise K Moyo, Mabaya Lucy, Mushiri Tawanda, Manasa Justen

## Abstract

The SARS-CoV-2 Omicron variant of concern (VOC) has multiple mutations in the spike (S) protein, which mediates viral infection and immunity. We analysed a sub-lineage of Omicron, designated XBB, that showed structural and functional changes in the S protein in response to the African selection pressures. We used molecular modelling to compare the S protein structures of Omicron and XBB and found that XBB had a reduced receptor-binding domain (RBD) due to the loss of some β-sheets, which may increase its affinity to the human angiotensin-converting enzyme 2 (hACE2) receptor. We also used Fast Unconstrained Bayesian AppRoximation (FUBAR) and Recombination Detection Program 4 (RDP 4) to perform selection and recombination analysis of the S protein sequences of Omicron and XBB and detected signals of positive selection and recombination in the N-terminal domain (NTD) of the S1 subunit, which contains antibody-binding epitopes, and the RBD, which is involved in viral entry. Our results reveal the structural and functional adaptation of the Omicron XBB variant in Africa and its potential implications for viral pathogenesis and immunity.

## Introduction

SARS-CoV-2 (severe acute respiratory syndrome coronavirus 2) is a novel coronavirus that emerged in late 2019 in China and caused a global pandemic of respiratory illness known as COVID-19 (coronavirus disease 2019) (Urhan, and Abeel, 2021). The SARS-CoV-2 virus causes lower respiratory tract infections and may lead to acute respiratory distress syndromes. The spread of SARS-CoV-2 and its negative effects around the globe have made it one of the most infamous pandemics ever documented in human history. The pandemic has resulted in more than 6.92 million fatalities among 776 million documented cases as of May 12, 2023 (WHO, 2023). Similar to other coronaviruses, the genetic material of SARS-CoV-2 comprises approximately 30 kilobases and encodes 29 proteins, including 16 nonstructural proteins, 9 accessory proteins, and 4 structural proteins (the spike protein, envelope protein, membrane protein, and nucleocapsid protein) (Jungreis *et al*., 2021; Bai *et al*.,2022).

Currently, the only variants of concern (VOCs) in circulation are Omicron XBB.1.9.2, which is a recombinant of omicron BA.2.10.1 and BA.2.75 sub-lineages. Conversely, Alpha (B.1.1.7), Beta (B.1.351), Gamma (P.1), and Delta (B.1.617.2) are now classified as previously circulating VOCs, (Dhama *et al*., 2022). The omicron variant was initially detected in South Africa and Botswana in November 2021 and consequently disseminated globally at a rapid pace, (Gowrisankar *et al*., 2022). According to preliminary research put forth by Uraki et al. (2023), it appears that omicron XBB (XBB) and BQ.1.1 entails a higher probability of reinfection when compared to the other currently circulating sub-lineages of omicron. Recent research by Miller et al. (2023) suggested that the XBB variants have the potential to reduce the efficacy of extant mRNA vaccines. The SARS-CoV-2 virus’s genetic evolution and adaptation is a growing concern for the COVID-19 pandemic. Evolution in viruses is primarily determined by the velocity at which mutations occur and disseminate across populations. Omicron’s high infectivity is likely due to its numerous mutations, containing over 60 substitutions/ deletions/ insertions - more than any other SARS-CoV-2 variant. The deletions and mutations facilitate enhanced transmissibility, augmented viral binding affinity, and elevated antibody evasion (Greaney *et al*., 2021).

It is suggested that the variant may have accrued mutations over time during its circulation within chronically infected individuals, particularly the immunocompromised with suboptimal ability to eliminate the viral infection (Kandeel *et al*., 2022). In September 2020, the Alpha variant (B.1.1.7) variant was isolated from a patient diagnosed with cancer in the United Kingdom. One year later, omicron BA.1 was isolated from an HIV-infected patient in Botswana and South Africa. Additionally, this shows that various types of immunocompromised states of the person may predispose to the selection of coronavirus variants (Tarcsai *et al*., 2022). The emergence of the omicron variant is not solely attributed to its immune evasive properties, but rather the alternative mechanisms of cellular entry that enable it to bypass the interferon-induced antiviral factors. Through the mutations unique to the omicron spike protein, the cellular entrance of omicron does not depend on serine transmembrane proteases but on matrix metalloproteinases, which facilitate membrane fusion. This entry pathway unlocked by the omicron spike enables evasion of interferon-induced factors that restrict SARS-CoV-2 entry following attachment. Therefore, the heightened transmissibility by the omicron is also due to its superior ability to invade nasal epithelia and its resistance to the cell-intrinsic barriers that are prominently present in the epithelia (Shi *et al*., 2023).

The salient characteristic of the XBB variant is its profound resistance to humoral immunity induced by vaccination against viral pathogens. A recent study by Tamura et al. (2023) demonstrated that 10 out of 14 breakthrough BA.2 infection sera and 9 out of 20 breakthrough BA.5 infection sera fail to neutralize XBB.1. They concluded that the collective effect of numerous substitutions present in the XBB.1 spike protein aids in promoting immune resistance of XBB.1. It was also observed that the immune-resistance exhibited by XBB.1 variant is strongly correlated to the Y144del mutation located in the N-terminal domain. The presence of six mutations (S373P, S375F, R408S, S447N, Q498R, and N501Y) on the S protein has been identified as being involved in the binding of ACE2. This discovery suggests that the presence of these modifications at the S protein level could enhance the ability of the viral protein to adhere to the ACE2 receptor, thus promoting host infection. (Zappa *et al*. 2023). A study by Daniloski *et al*. (2021) showed that the substitution of aspartate with glycine at the D614G mutation in the S gene leads to higher transduction rates in human epithelial cells. A recent study has discovered that the R203K and G204R mutations in the N gene are responsible for enhancing viral fitness (Leary *et al*. (2020).

Besides genetic mutations, the evolutionary trajectory of the novel coronavirus SARS-CoV-2 can be attributed to the phenomenon of genetic recombination. Recombination represents a mechanism that can accelerate the process of viral adaptation by facilitating the creation of hybrid variants by combining mutations from distinct genetic backgrounds (Markov *et al*. 2023). Recombination requires simultaneous co-infection of a host with two genetically divergent viruses, whose recombination results in viable offspring that disseminate to other hosts.

The Omicron XBB variant is a result of the recombination of BA.2.10.1 and BA.2.75 sub-lineages. The comparative analysis of genomic sequences across various coronaviruses offers valuable insights into the identification and regions that have undergone positive selection during the course of their individual evolutionary trajectories. Currently, there are large SARS-CoV-2 genome sequence datasets that scientists can use to monitor the evolutionary patterns of SARS-CoV-2 and ascertain whether any particular sites present indications of adaptive evolution. SARS-CoV-2 evolution can be influenced by viral and host factors. Host factors, including the host species (animal or human), viral shedding time, and immune system status, have an impact on the diversity of SARS-CoV-2 (Gandi *et al*., 2022; Voloch *et al*., 2021). Prolonged infections in individuals with compromised immune systems have the potential to expedite viral evolutionary processes, which may result in the emergence of distinct variants (Sonnleitner *et al*., 2022).

Little is known about the evolution and adaptation of the XBB variant in different geographic and epidemiological contexts of Africa. In particular, there is a lack of data and analysis on the mutations being acquired by the omicron sub-lineages circulating in Africa, where the variant originated, and where diverse environmental and host factors may exert different selection pressures on the virus. This research project aims to identify and characterize the mutations being acquired by SARS-CoV-2 omicron sub-lineage XBB in Africa due to the selection pressures specific to Africa. This research will provide insights into this sub-lineage’s evolutionary dynamics and adaptive potential and its implications for disease transmission, severity, immunity, and intervention strategies. In this research project, we collected genomic data of SARS-CoV-2 omicron sub-lineage XBB from various African countries and regions, using publicly available database Global Initiative on Sharing All Influenza Data (GISAID) (Khare *et al*., 2021). Analysis was done in 3 parts namely selection analysis, recombination analysis and 3D structure alignment against the reference SARS-CoV-2 omicron XBB 3D structure.

This article is divided into three parts: 1) Selection Analysis, 2) Recombination Analysis, and 3) 3D-Structure Prediction and Comparison. Each section includes methodology, results, and discussion combined into one cohesive piece.

### 1. Selection Analysis

Several Python libraries were used to load, manipulate and visualise data in this project. Pandas was used to load (McKinney, 2010).csv files into DataFrames/Tables, Biopython (Cock *et al*.,2009) was used to interact with biological data using Python language, and matplotlib (Hunter, 2007) and seaborn (Waskom *et al*., 2021) were used to visualize data into plots or graphs.

The spike gene region was extracted from the whole genome sequences of SARS-CoV-2 using the nucleotide position range of 21 500 to 25 500, which is an approximation to accommodate the different lengths of the sequences. The spike gene locus in the reference genome is from 21 563 to 25 384. Duplicate sequences and sequences containing more than four unidentified nucleotide bases (represented by N) were removed from the dataset to reduce noise in the downstream analysis.

Multiple sequence alignment was performed using Clustal O (Sievers *et al*., 2011) hosted on the European Bioinformatics Institute (EBI) webserver, with the following parameters: guidetreeout: true, dismatout: false, dealign: false, mbed: true, mbediteration: true, iterations: 0, gtiterations: −1, hmmiterations: −1, outfmt: fasta, order: aligned, stype: dna. The alignments were edited with MEGA X software (Stecher., 2020), using the reference spike gene sequence from GISAID to determine the accurate location of each spike gene within the approximate gene region.

A phylogenetic tree was generated from the aligned sequences using MEGA X software with the Maximum Likelihood method and bootstrap number = 100. The phylogenetic tree was exported in newick format for two purposes: (1) to visualise the relationship between sequences and their evolutionary distances using interactive tree of life (iToL) (Letunic and Bork, 2016), which generated an unrooted phylogenetic tree (shown below), and (2) to perform selection analysis using Fast Unconstrained Bayesian AppRoximation (FUBAR) (Murrell *et al*. 2013) on sequences above 500, which required a guide tree to be integrated with the sequence data. NCBI G-Workbench was used to integrate the guide tree into the spike sequence file.

FUBAR pervasive selection analysis tool hosted on the Datamonkey (Weaver *et al*., 2018) webserver was used to investigate sites undergoing both positive and negative pervasive selection. The parameters for FUBAR analysis were set as follows: number of grid points = 20, concentration parameter of the Dirichlet prior = 0.5. FUBAR results were exported in table of selection results for each site in .csv format below. Sites under pervasive positive or negative selection were identified by applying a posterior probability cutoff of 0.95 for β>α or β<α, respectively.Containing two domains that are critical for viral infection and immunity, the S1 subunit of the SARS-CoV-2 spike (S) protein has acquired numerous mutations in these domains, which may affect its interaction with the human angiotensin-converting enzyme 2 (hACE2) receptor and the host antibody response.

**Figure 1:**
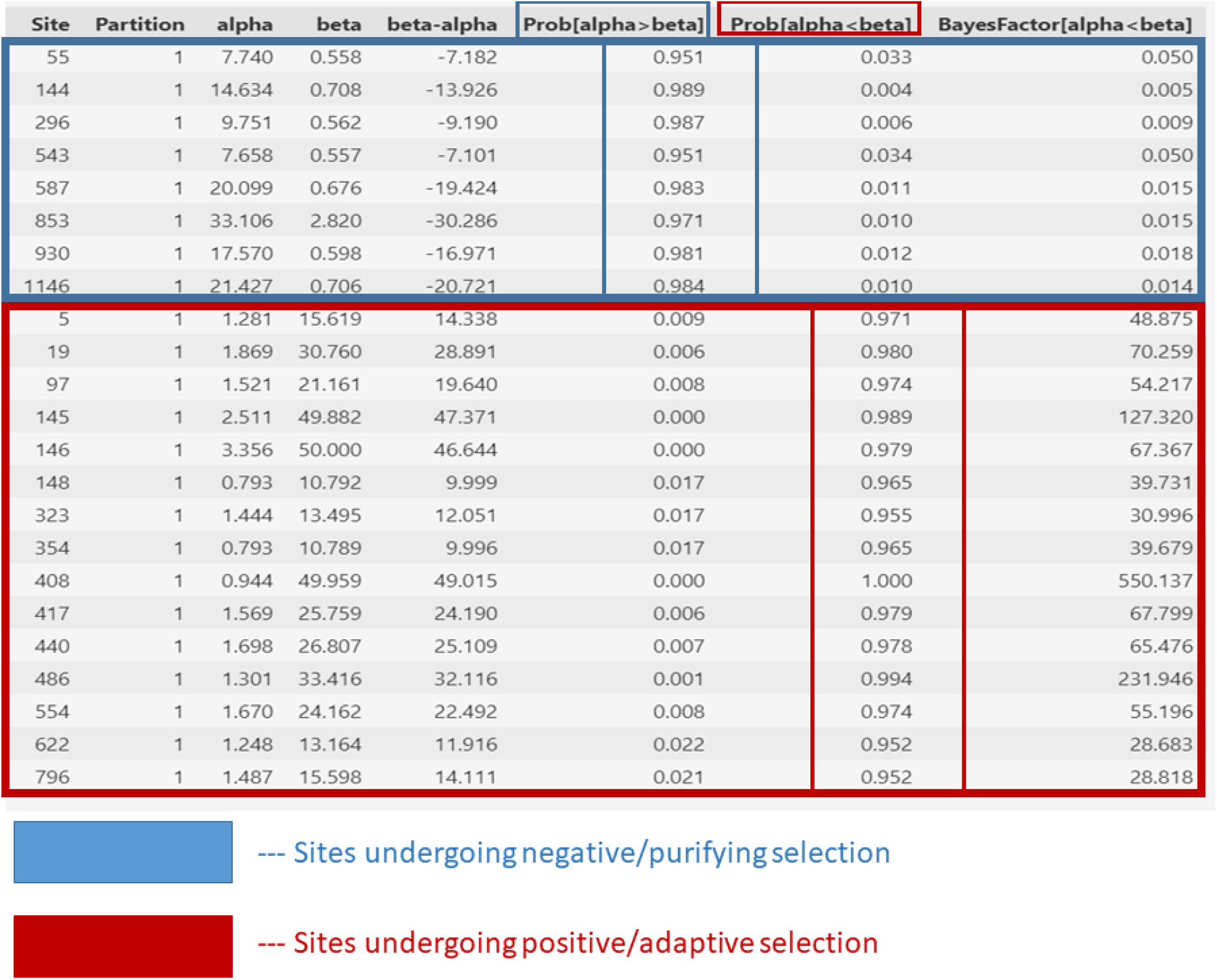
FUBAR analysis results for sites under per vasive selection. Sites under strong purifying selection (β<α) are highlighted in blue, and sites under strong adaptive selection (β>α) are highlighted in red. The parameters for each site are shown in the corresponding columns.

**Figure 2:**
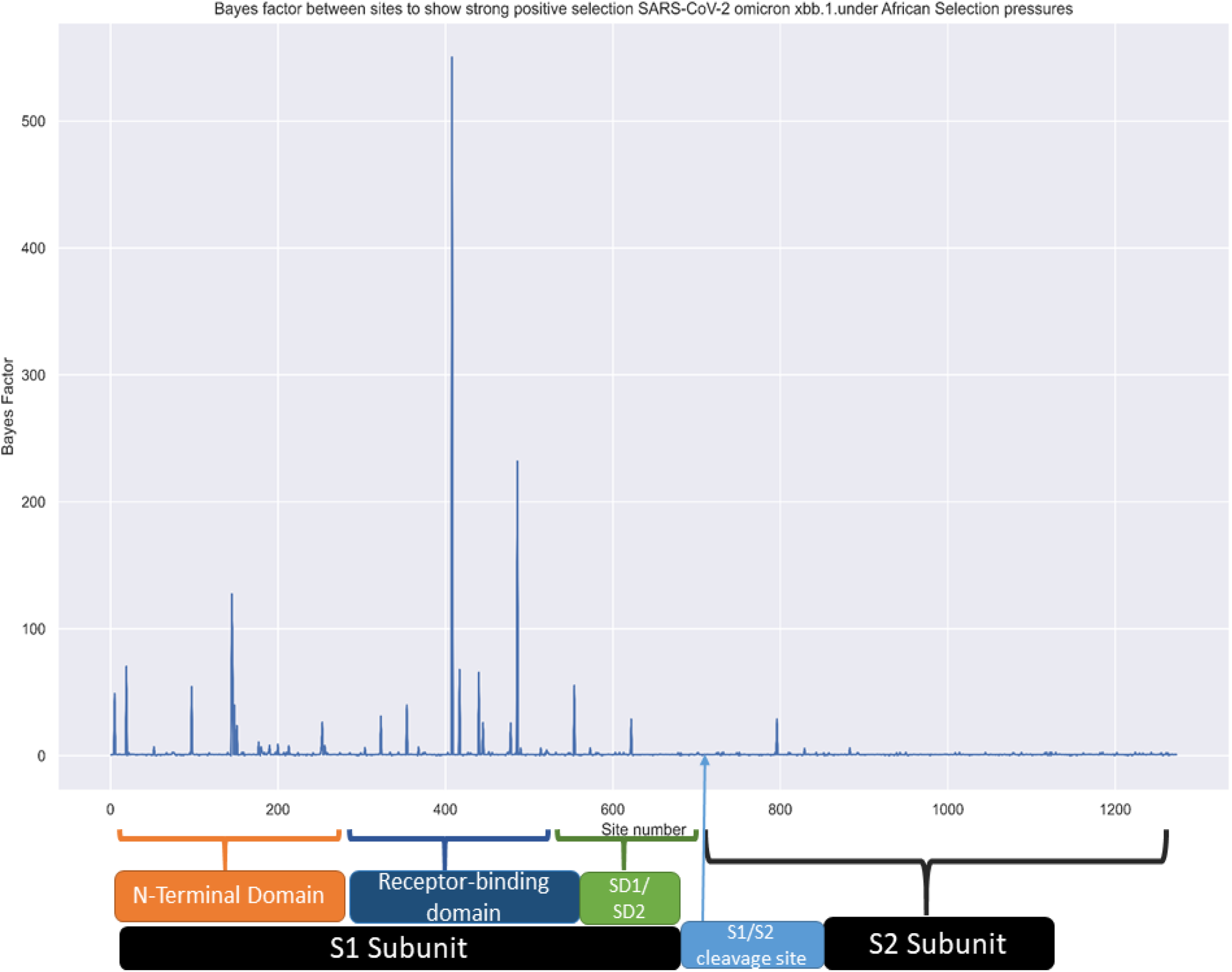
Bayes Factor (BF) for pervasive positive selection at each site from FUBAR results. BF was used to test the hypothesis of β > α at each site, where a higher BF indicates stronger support for positive selection. The bar plot shows the sites with high BF values as peaks, indicating evidence of pervasive positive selection.

Performing a pervasive selection analysis of the S1 subunit sequences of Omicron from Africa using Fast Unconstrained Bayesian AppRoximation (FUBAR), this study identified sites under strong positive or negative selection and investigated their functional implications. Finding that the receptor-binding domain (RBD) had the highest number of sites under positive selection, followed by the N-terminal domain (NTD), both involved in enhancing viral infectivity and escape from pre-existing immunity respectively, positive selection in this domain may also require more mutations for optimal viral entry in the highly diverse African gene pool.

Among the RBD mutations, high levels of positive selection were observed at amino acid positions 408 and 486, while the D614G mutation, which was previously associated with increased transmissibility, showed reduced selection pressure.

Detecting strong positive selection at site 145, which is a known target of neutralizing antibodies, this study also found that positive selection in the NTD may confer immune evasion and delay antibody recognition. Providing insights into the molecular evolution and adaptation of the Omicron variant of concern (VOC) in Africa and its potential impact on viral pathogenesis and immunity, this study highlighted the importance of monitoring the S1 subunit mutations of SARS-CoV-2.

### 2. Recombination Analysis

The SISCAN method (Sun *et al*., 2021) implemented in the recombination detection program (RDP 4) (Martin *et al*., 2015) was used to detect recombination events among SARS-CoV-2 S sequences. This method is a Monte Carlo procedure that measures variations in phylogenetic signals in gene sequences that result from recombination. It scans a window of fixed size along the alignment and calculates the Z-score of the phylogenetic signal for each sequence at each position. The Z-score is a measure of how many standard deviations a value is away from the mean. A high Z-score indicates a strong phylogenetic signal, while a low Z-score indicates a weak or contradictory phylogenetic signal. The method then compares the Z-scores of the potential recombinant sequence with those of the parental and outlier sequences to identify regions where the phylogenetic signal is inconsistent with the expected topology.

The SISCAN method requires a minimum of three or four sequences for each analysis: a parental sequence, a potential recombinant sequence and an outlier sequence. The following sequences were used for the analysis:

- A parental sequence from a Southern African omicron study (Vianna *et al*. 2022)
- A potential recombinant sequence derived from a consensus of African S sequences.
- An outlier sequence from a GISAID S wild type sequence.

**Figure 3:**
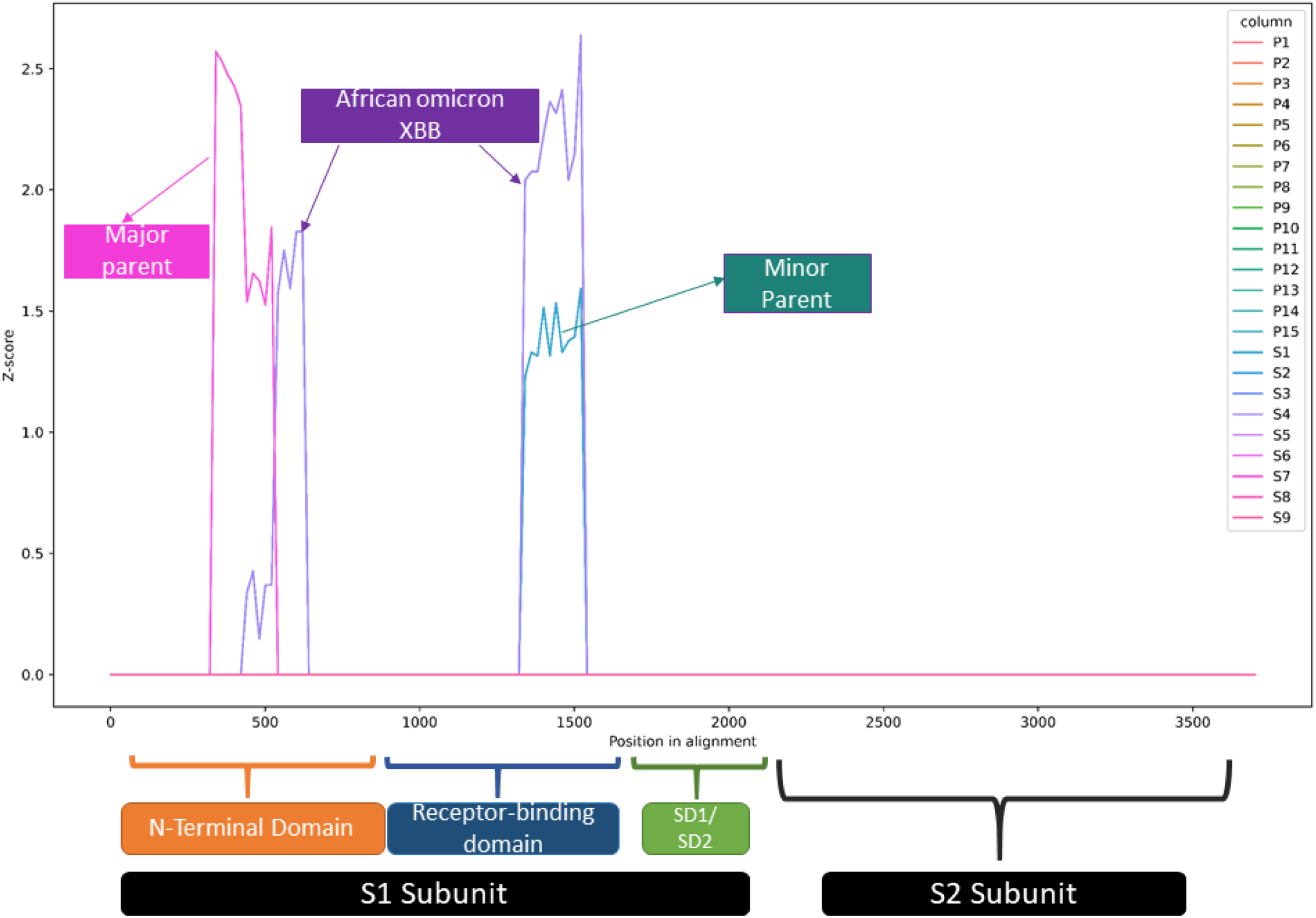
Z-score plot of the phylogenetic signal for each sequence at each nucleotide position from SISCAN analysis. The purple line represents the African Omicron XBB sequence, which is the potential recombinant. The pink line represents the GISAID S gene reference sequence, which is the major parent. The green line represents the Southern Africa Omicron S sequence, which is the minor parent. The plot shows regions of inconsistent phylogenetic signal between the potential recombinant and the parents, indicating recombination events. These regions are around nucleotide number 400-600 (amino acid number ∼ 133-200 of the spike protein), which is in the N-terminal domain (NTD), and around nucleotide number 1350-1550 (amino acid number ∼ 450-500 of the spike protein), which is in the receptor binding domain (RBD).

SISCAN is a powerful tool that can identify both inter- and intra-species recombination, and multiple breakpoints, and distinguish between compositional and recombination signals. However, it may also have some drawbacks, such as being dependent on window size, alignment quality, and sequence divergence.

Evidence of recombination was found in the N-terminal domain (NTD) and the receptor-binding domain (RBD) regions of the S protein. The major parent of the recombinant omicron XBB was the SARS-CoV-2 wild type, while the minor parent was the omicron variant from a Southern African study by Vianna et al. 2022.

The NTD region, especially around residues 133-200, has been shown to undergo reversions that disrupt the binding of NTD-specific neutralizing antibodies and increase the viral load (Shen *et al*., 2021). This may explain why the authors detected recombination signals in this region, as the omicron XBB variant reverted to its ancestral state.

The RBD region, which is critical for viral entry and transmissibility, also showed recombination signals between omicron XBB and omicron variants. This may have resulted from the co-infection of a host with both variants, leading to the exchange of RBD fragments. The omicron XBB variant may have gained an adaptive advantage from acquiring mutations from the omicron variant, which has high fitness in the African population.

### 3. 3D protein structure alignment and analysis

Jalview 2.11 (Waterhouse *et al*., 2009) was used to translate the nucleotide alignment of the spike gene sequences from different SARS-CoV-2 variants into amino acid sequences and to infer the consensus sequence for each variant. The protein structure of the African Omicron XBB variant was predicted using the PHYRE 2.0 (Kelley *et al*.,2015) web server, with the crystal structure of the Omicron spike protein (pdb id = 7T9K) as a template. The resulting 3D model of the African Omicron XBB spike protein had 100% accuracy and high confidence.

Structural alignment of the African Omicron XBB spike protein with the Omicron spike protein (pdb id = 7T9K) was performed to compare their structural features. The figure below shows the superimposed structures of the two proteins, with the Omicron spike protein in blue and the African Omicron XBB S protein in pink ribbons. The African Omicron XBB S protein was smaller than the Omicron spike protein, which could affect its binding to host receptors and antibodies.

**Figure 4:**
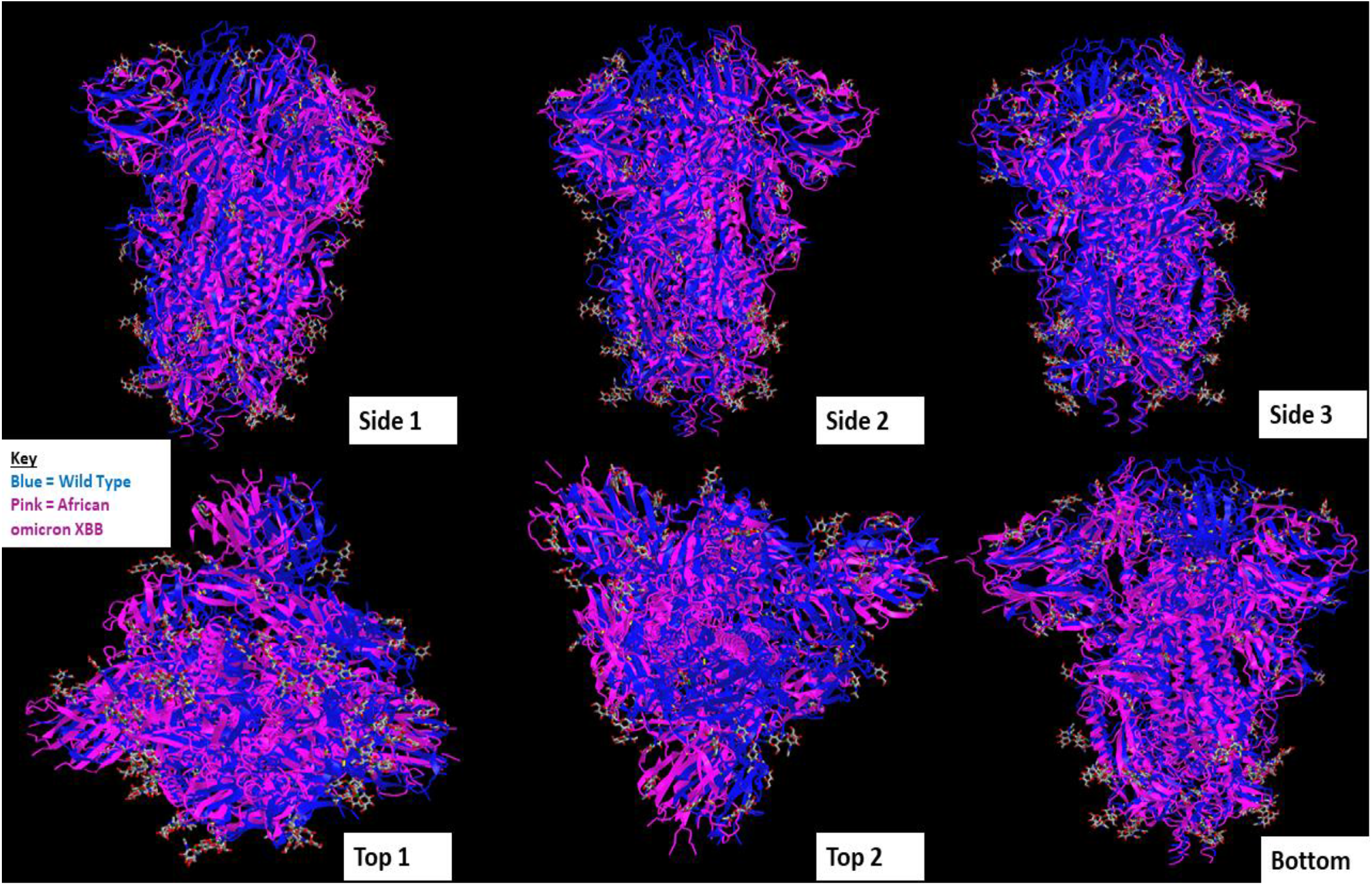
Structural alignment of the omicron spike protein (blue) and the African Omicron XBB S protein (pink) in three different views: side, top and bottom.

To examine the structural compactness of the RBD, we used JPred (Bryson and Jones, 2012) in Jalview to predict the secondary structures, as shown in Figure 5.

**Figure 5:**
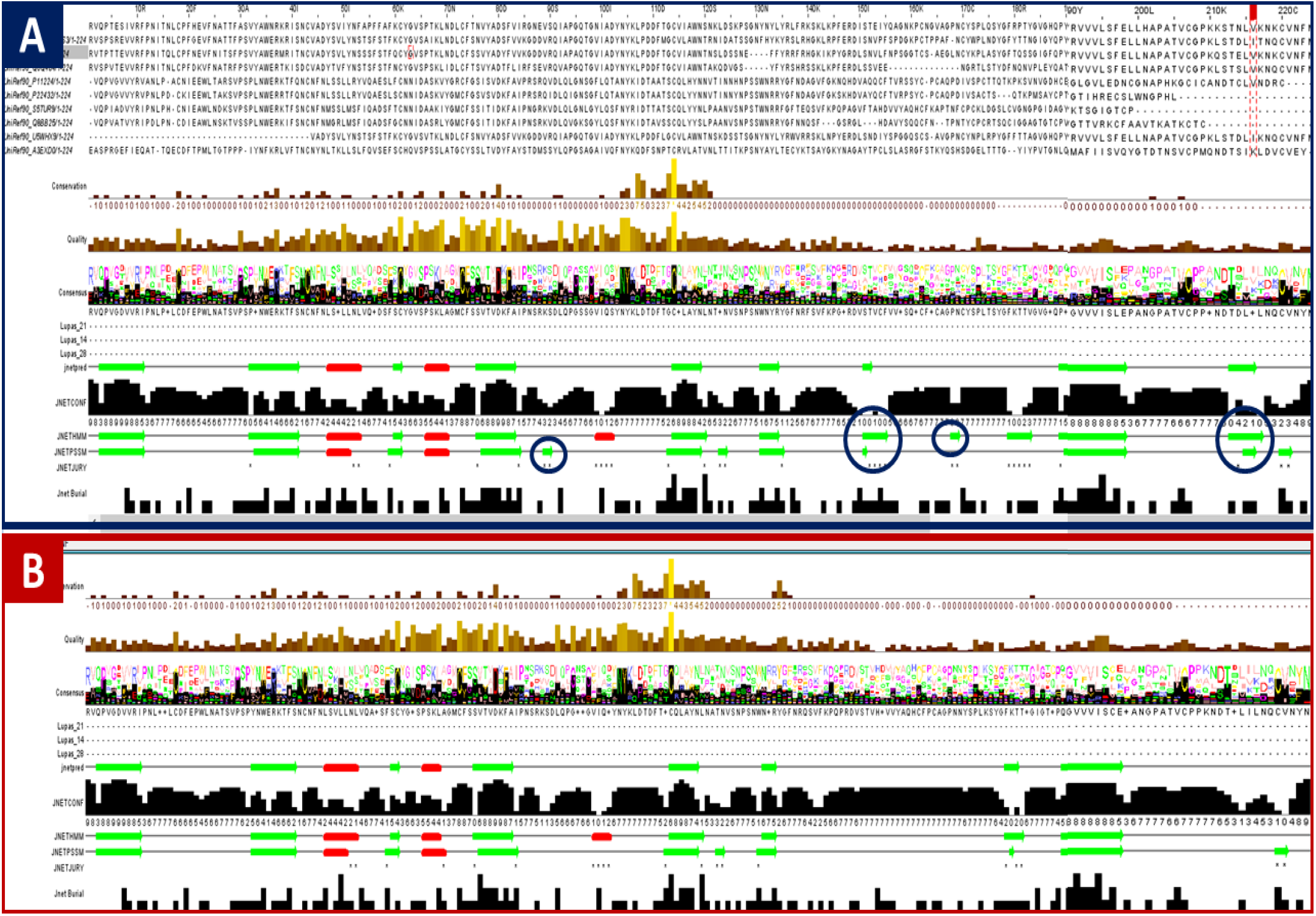
Structural differences between the RBDs of the SARS-CoV-2 Southern African omicron and the omicron XBB variant from the African population. The β-sheets that are present in the Southern African omicron RBD (A) but deleted in the omicron XBB RBD (B) are highlighted in blue.

Comparing the structure of the receptor-binding domain (RBD) of the SARS-CoV-2 omicron XBB spike (S) protein from the African population with that of the omicron variant, the authors observed that the omicron XBB RBD had four β-sheet deletions at residues V407-S408, S469-Y473, P486-N487 and T531-K535, relative to the omicron RBD. These deletions resulted in a smaller RBD size, which may enhance its affinity to the human angiotensin-converting enzyme 2 (hACE2) receptor and increase the viral entry and transmissibility of the omicron XBB variant.

The authors also hypothesized that the smaller RBD size may be an adaptation to the diverse hACE2 phenotypes in the African population, which may impose selective pressure on the virus to optimize its entry efficiency.

## Conclusion

The African omicron XBB variant exhibited structural and functional changes as an adaptation to the African selection pressures in the receptor-binding domain (RBD) and N-terminal domain (NTD) of the SARS-CoV-2 spike (S) protein. The omicron XBB S protein was observed to be smaller than the omicron S protein due to the absence of some β-sheets in the RBD, which might enhance its fitness to the human angiotensin-converting enzyme 2 (hACE2) receptor. Enhanced fitness may increase the successful interaction between the S protein and the hACE2 receptor, facilitating viral entry and ensuring survival and transmissibility of the XBB variant. Selection and recombination signals in the NTD of the S1 subunit of the omicron XBB S protein were also detected in the analysis. The NTD is a site for antibody-binding epitopes, and the XBB variant may be evolving in this region to evade the host immune response, which enhances its survivability and reproduction rates.

## Acknowledgment

The study was not funded.

